# Integrative analysis identified common and unique molecular signatures in hepatobiliary cancers

**DOI:** 10.1101/2021.10.06.463304

**Authors:** Nabanita Roy, Ria Lodh, Anupam Sarma, Dhruba Kumar Bhattacharyya, Pankaj Barah

**Affiliations:** Department of Molecular Biology and Biotechnology, Tezpur University, Napaam, Sonitpur, Assam, 784028, India; Department of Oncopathology, Dr. Bhubaneswar Borooah Cancer Institute, Guwahati, Assam, 781016, India; Department of Computer Science and Engineering, Tezpur University, Napaam, Sonitpur, Assam, 784028, India

**Keywords:** Hepatobiliary cancers, Transcriptomics, Systems biology, Biological networks, Hub genes, Systems biomarker

## Abstract

Hepatobiliary cancers (HBCs) are the most aggressive and sixth most diagnosed cancers globally. Biomarkers for timely diagnosis and targeted therapy in HBCs are still limited. Considering the gap, our objective is to identify unique and overlapping molecular signatures associated with HBCs. We analyzed publicly available transcriptomic datasets on Gallbladder cancer (GBC), Hepatocellular carcinoma (HCC), and Intrahepatic cholangiocarcinoma (ICC) to identify potential biomarkers using integrative systems approaches. An effective *Common and Unique Molecular Signature Identification (CUMSI)* approach has been developed, which contains analysis of differential gene expression (DEG), gene co-expression networks (GCN), and protein-protein interactions (PPIs) networks. Functional analysis of the DEGs unique for GBC, HCC, and ICC indicated that GBC is associated with cellular processes, HCC is associated with immune signaling pathways, and ICC is associated with lipid metabolic pathways. Our findings shows that the hub genes and pathways identified for each individual cancer type of the HBS are related with the primary function of each organ and each cancer exhibit unique expression patterns despite being part of the same organ system.

## 1. Introduction

Hepatobiliary cancer (HBC) refers to cancers of the hepatobiliary systems, which comprises of hepatocellular carcinoma (HCC), gallbladder cancer (GBC), and cholangiocarcinoma (CCA). CCA further includes intrahepatic cholangiocarcinoma (ICC), and extrahepatic cholangiocarcinoma (ECC). GBC and CCAs are collectively referred to as biliary tract cancers (BTC)(A. Bailey & Shah, 2019; Shibata et al., 2018). HCC is the seventh most common cause of cancer-related death, while GBC is the most aggressive and common malignancy of the BTCs (Brägelmann et al., 2021; Nepal et al., 2021; Xue et al., 2020). The incidence rates of HCC are higher in East Asian countries whereas, BTCs are more frequent in Southeast Asian countries (Shibata et al., 2018). The epidemiological status for each cancer of the hepatobiliary tract is also specific. HCC development is largely associated with alcoholism, hepatitis and metabolic diseases such as obesity, whereas CCAs are mostly linked with bile duct anomaly and sclerosing cholangitis (Giudicessi & Ackerman, 2013; Yang et al., 2019). The well-known risk factor associated with GBC is gallstone disease (Lazcano-Ponce et al., 2001).

The International Cancer Genome Consortium (ICGC) has conducted OMICs based analysis on HCC cases from different epidemiological backgrounds and ethnic populations. Currently genomic and transcriptomic datasets on more than 1000 HCC cases are publicly available. The key driver genes identified from these genomic studies on HCC are TP53, CTNNB1 and TERT (Craig et al., 2020). The TERT promoter mutations (TPMs) are significantly linked with HCC development (Nault et al., 2014; Shibata et al., 2018; Totoki et al., 2014). The analysis of molecular data from CCA cases also identifies a few driver genes which include TP53, ARIDIA, KRAS, SMAD4, PTEN, and ERBB2 (Jusakul et al., 2017). However, compared to that of HCC and CCA, there are very limited genomic as well as transcriptomic data on GBC. From whole exome sequencing, it has been observed that TP53, KRAS and ERBB3 are the most potent driver genes in GBC. The list of potential driver genes is almost similar in all the cancers of the hepatobiliary tract except for TPM which is highly mutated in HCC (Nakamura et al., 2015). Since the characteristics of individual cancer and tumor microenvironment are different, it is therefore important to investigate the overlapping and specific molecular signatures and their associated cellular/metabolic pathways in HBCs to develop targeted therapy for each cancer type of the hepatobiliary tract. An integrative systems biology approach for studying hepatobiliary system enables the understanding of biological functions of the liver and its interaction with other tissues all owing us to build biological network models such as metabolic, transcriptional regulatory, protein–protein interactions, signaling and co-expression networks (Teufel, 2015). Such comprehensive systems level analysis may shed some light into the dynamics of the hepatobiliary system in a holistic manner and elucidate the molecular mechanisms involved in disease progression (Mardinoglu et al., 2018). As most genes involved in disease pathology are co-regulated, studying gene expression across the whole genome via an omics approach helps in the identification of the group of genes that are co-regulated during cancer. Characterization of the entire genetic landscape through transcriptome sequencing can help find biomarkers for early identification of high-risk individuals, prompt diagnosis, prognosis, prediction, and therapeutic targets for HBC patients (Teufel et al., 2012).

We have developed an effective **C**ommon and **U**nique **M**olecular **S**ignature **I**dentification (CUMSI) approach for studying hepatobiliary cancers. CUMSI approach enables us to analyze publicly available transcriptomic datasets using benchmarked computational pipelines to identify overlapping and unique DEGs in cancers of the HBS. The significant DEGs identified have been further analyzed via computational approaches that include ensemble approach for differential gene expression analysis, gene co-expression network analysis, protein-protein network analysis and transcriptional regulatory network analysis to identify systems level biomarkers for each selected cancer type of the hepatobiliary tract. The conceptual workflow has been indicated in **Figure 1**.

**Figure 1:**
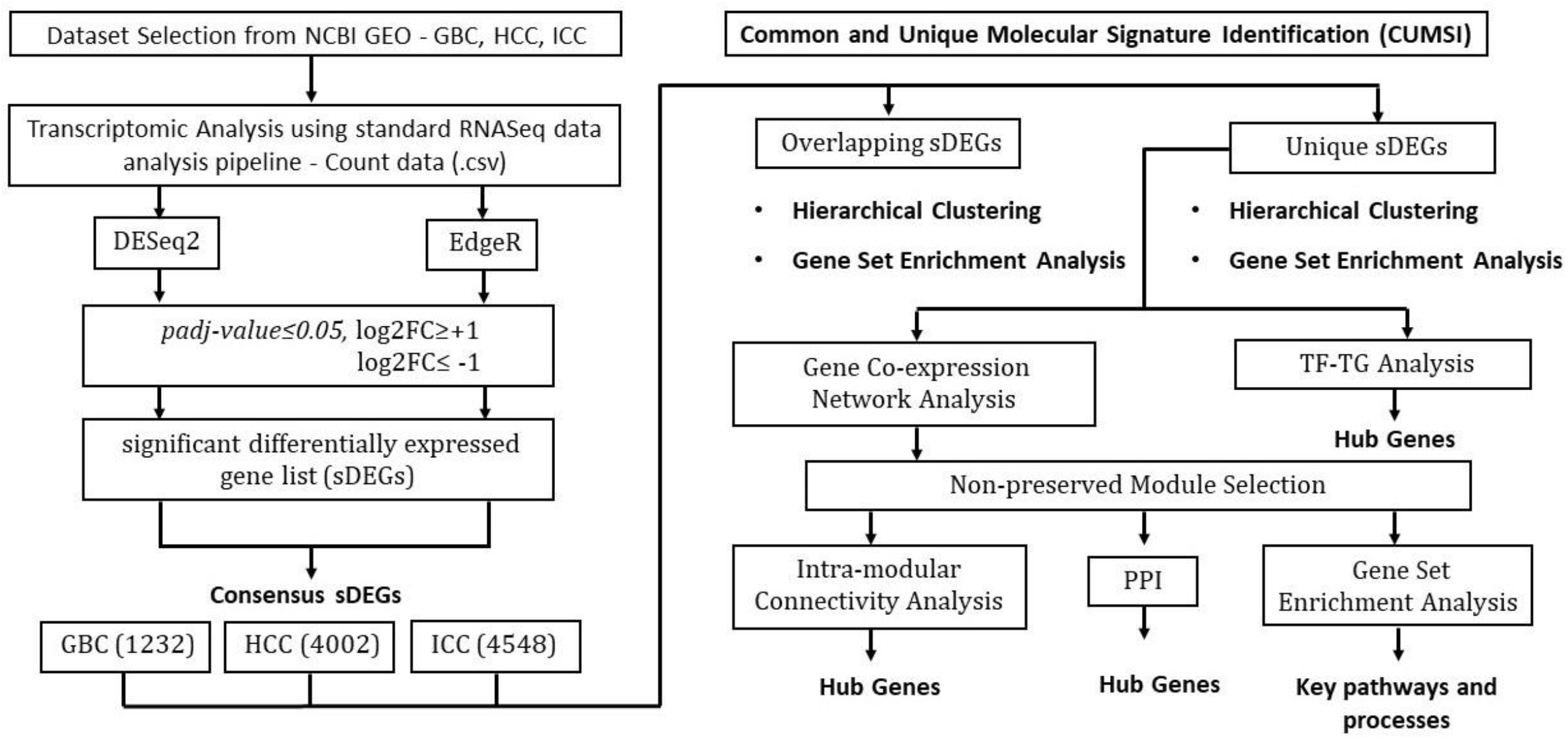
Schematic workflow of the study to find systems biomarker for hepatobiliary system cancers using CUMSI approach.

## 2. Results

### 2.1 The overlapping and unique gene expression profiles of GBC, HCC, and ICC

For identifying the overlapping and specific DEGs among GBC, HCC & ICC, differential gene expression analysis has been carried out using two algorithms. For each cancer dataset, the consensus of both DESeq2 and EdgeR is taken for screening of significant DEGs (sDEGs) (**Supplementary Figure S1**). We identified 256 overlapping DEGs among the three cancers, and 561, 2005, 2580 unique DEGs in GBC, HCC and ICC, respectively. Hierarchical clustering of the overlapping DEGs shows distinct gene expression patterns in each cancer and is mostly associated with cell cycle processes (**Supplementary Figure S2**). Moreover, the expression levels of the unique as well as the overlapping DEGs in GBC are significantly downregulated as compared to HCC and ICC, which suggested that GBC exhibits a distinct trend of gene expression pattern. Our cluster analysis on gene expression data for the top 500 unique sDEGs for all the three cancer types using complete linkage hierarchical approach identified distinct clusters for each cancer type as compared to normal samples. These distinctions in terms of co-expressed groups of genes are a clear indication that there exists a unique set of molecular signatures. These observations indicate that despite belonging to a common organ system, progression of GBC, HCC, and ICC results from differential patterns of gene expression. The overall unique and specific gene expression profiles are represented in **Figure 2**.

**Figure 2:**
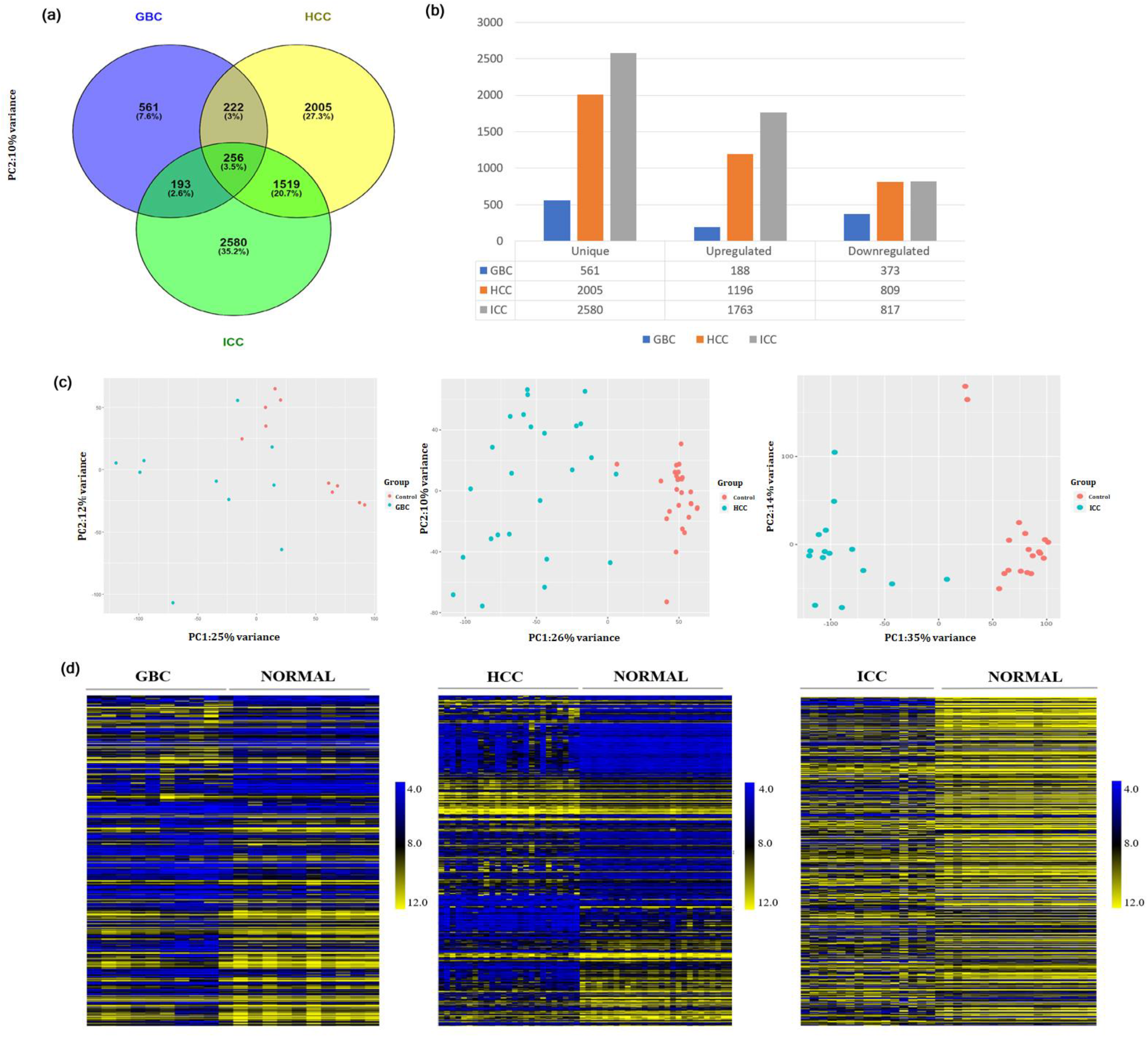
Data visualization. **(a)** Venn diagram constructed using sDEGs (*padj-value≤0*.*05*, log2FC±1) identified byDESeq2+EdgeR consensus. **(b)** Upregulated (log2FC≥1) and downregulated (log2FC≤-1) sDEGs of GBC, HCC, & ICC. **(c)** Principle component analysis of cancer and control samples of GBC, HCC, &ICC. **(d)** Heatmap constructed using sample expression values of top 500 unique sDEGs of GBC, HCC, and ICC.

### 2.2 Functional annotation and KEGG pathway analysis of unique DEGs in GBC, HCC, and ICC

To understand the association of unique DEGs in GBC, HCC, and ICC progression, the significant (*p-value≤0*.*05*) biological processes and KEGG pathways have been identified. Functional enrichment analysis of unique DEGs shows that cancers of the hepatobiliary tract are associated with distinct biological processes and cellular pathways, as given in **Supplementary Figure 3**. GBC is mainly linked with cellular processes. HCC is largely associated with immune system regulation and signaling pathways, whereas, ICC is associated with metabolic pathways, mostly lipid and steroid metabolism.

### 2.3 Identification of module eigengene and detection of non-preserved modules

For the construction of gene co-expression network (GCN) for each type of hepatobiliary cancer, the unique significant DEGs (561, 2005, and 2580) in GBC, HCC, and ICC have been taken as input. GCNs for each cancer type are constructed using WGCNA package in R and an adjacency matrix has been built by raising the Pearson’s correlation coefficient of each pair of DEGs to the soft thresholding power β for satisfying a scale-free co-expression network. The soft-thresholding power β for (i) GBC and control are 12 and 9, respectively, for (ii) HCC and control 12, and for (iii) ICC and control 14 and 12; respectively (**Supplementary Figure S4**). Using hierarchical clustering, modules for cancer and control networks have been identified. Distances among the modules are calculated using dissimilarity between module eigengenes (i.e., representative of the gene expression profiles in a module), and similar modules are combined. A total of 5, 10 and 23 modules are identified from GBC, HCC, and ICC network, respectively. In the module preservation analysis for each cancer type to its respective control, the brown, black, and lightyellow modules have been identified as non-preserved, having a *Z-summary*- 0.87, 0.12, 0.02 and *medianRank*- 4, 10, 23 respectively, as represented in **Figure 3** and **Supplementary Table S2**.

**Figure 3:**
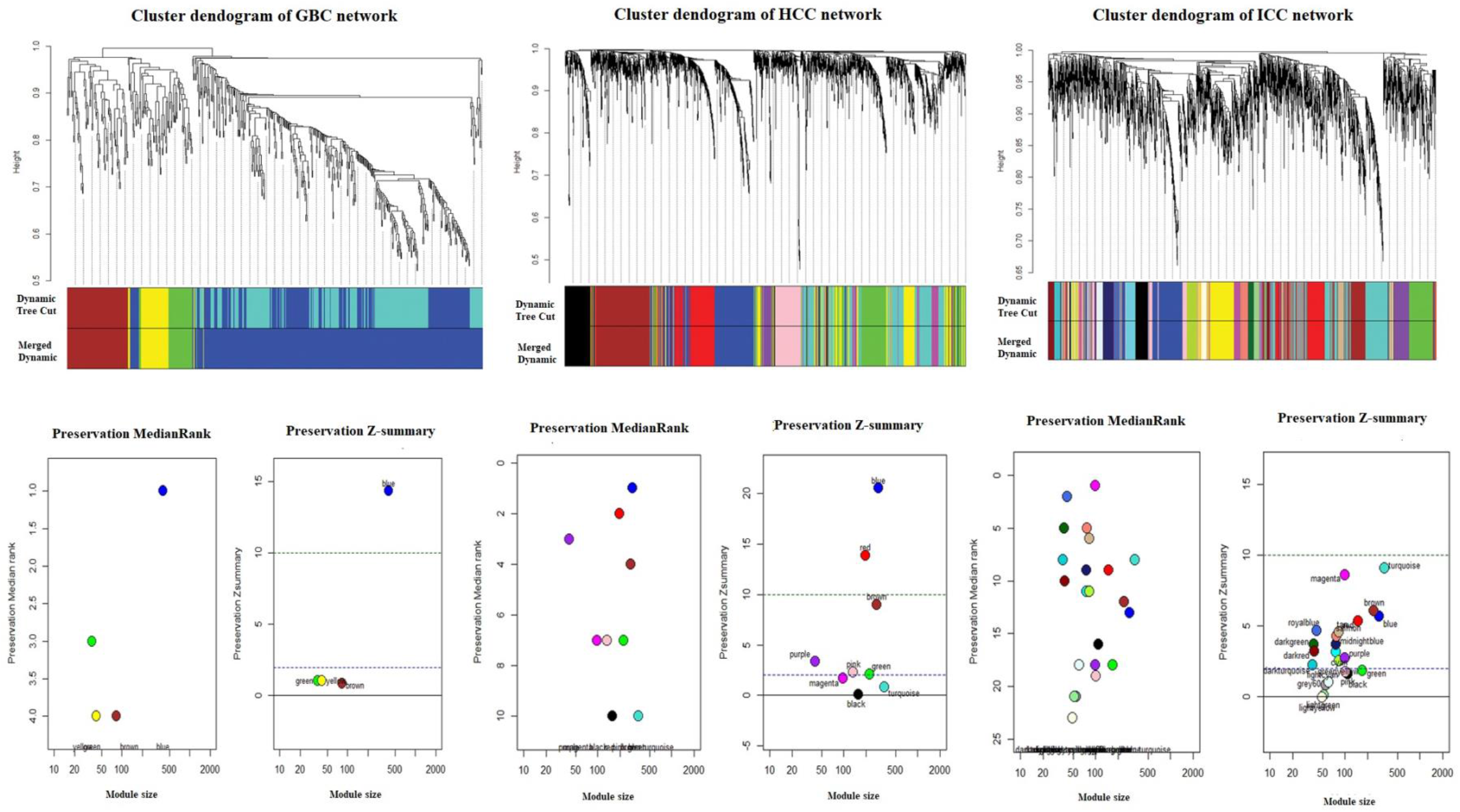
Gene co-expression network and module preservation analysis. **(a)** Clustering dendrogram of genes in GBC, HCC, & ICC network based ondistances between gene pairs that are subsequently grouped into modules (minClusterSize=30) based on the similarity of the magnitude of gene expression. **(b)** Preservation analysis of modules based on *Z-summary* and *medianRank*.Modules whose topological properties changed in a cancer network compared to normal networks are termed non-preserved modules. The black, brown, and lightyellow modules are non-preserved in GBC, HCC, and ICC respectively.

### 2.4 Hub gene identification from non-preserved modules

The genes having a large number of interacting nodes in the non-preserved modules are recognized as hub genes using intra-modular connectivity analysis. The top five genes from each cancer type, having the highest correlation weight are considered as hubs. The weight of the potential candidate genes identified is indicated in **Table 1**. The hub genes identified for GBC, HCC, and ICC through intra-modular connectivity analysis are MAP3K13, AC069287.1 and TRPC1.

**Table 1:** Identification of hub genes from non-preserved modules through Intra-modular connectivity analysis and PPI network respectively.

| Intra-modular Connectivity |  |  |  |  |  |
| --- | --- | --- | --- | --- | --- |
| GBC |  | HCC |  | ICC |  |
| Gene | Weight | Gene | Weight | Gene | Weight |
| MAP3K13 | 6.76 | AC069287.1 | 2.24 | TRPC1 | 1.21 |
| HYLS1 | 5.71 | SEPTIN14P19 | 2.16 | SLC45A1 | 0.99 |
| KIAA0895 | 5.67 | SEPTIN14P5 | 1.75 | PPP2R3A | 0.92 |
| AC130456.2 | 5.51 | DCDC1 | 1.59 | BACE1 | 0.90 |
| GPR39 | 5.48 | CICP6 | 1.25 | SNPH | 0.90 |
| PPI |  |  |  |  |  |
| GBC |  | HCC |  | ICC |  |
| Gene | Degree | Gene | Degree | Gene | Degree |
| ERBB2 | 119 | ACTN2 | 71 | GLI1 | 69 |
| JUP | 67 | RFC3 | 49 | WWTR1 | 26 |
| EFNA1 | 37 | AXIN2 | 35 | VAV3 | 19 |
| AHR | 25 | UBQLN4 | 34 | **BACE1** | 15 |
| CD2AP | 23 | RAP2A | 26 | **TRPC1** | 14 |

PPI networks are constructed using the genes of the non-preserved modules for predicting the interactions of genes at the protein level from STRING database with confidence score> 0.4. The genes having the highest degree are identified as hubs and interactive PPI networks generated with DEGs of the nonpreserved modules are represented in **Figure 4**. The hub genes for GBC, HCC, and ICC are ERBB2, ACTN2, and GLI1, respectively and have been tabulated in **Table 1**.

**Figure 4:**
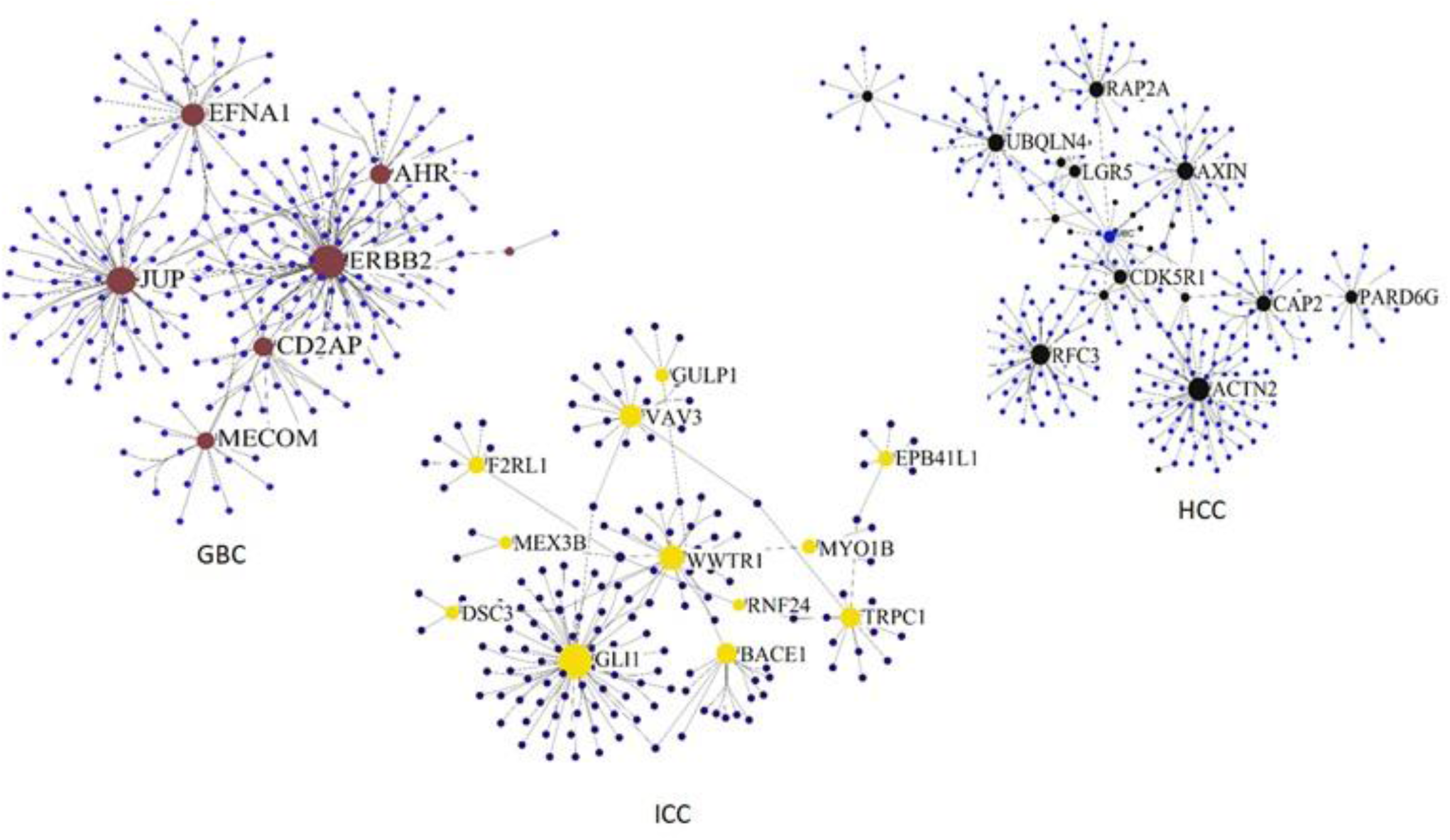
PPI network of the non-preserved modules in GBC, HCC, and ICC. The large brown, black, and yellow nodes represent the hub proteins for GBC, HCC, and ICC respectively that have the highest degree. Significant protein-protein interactions were filtered using a combined score > 0.7.

### 2.5 Functional annotation and pathways associated with genes of the non-preserved modules

The biological processes and pathways associated with the non-preserved modules (having a *p-value≤0*.*05*) of GBC, HCC, and ICC, are determined using the R package ClusterProfiler, and have been tabulated in **Table2**.

**Table 2:**
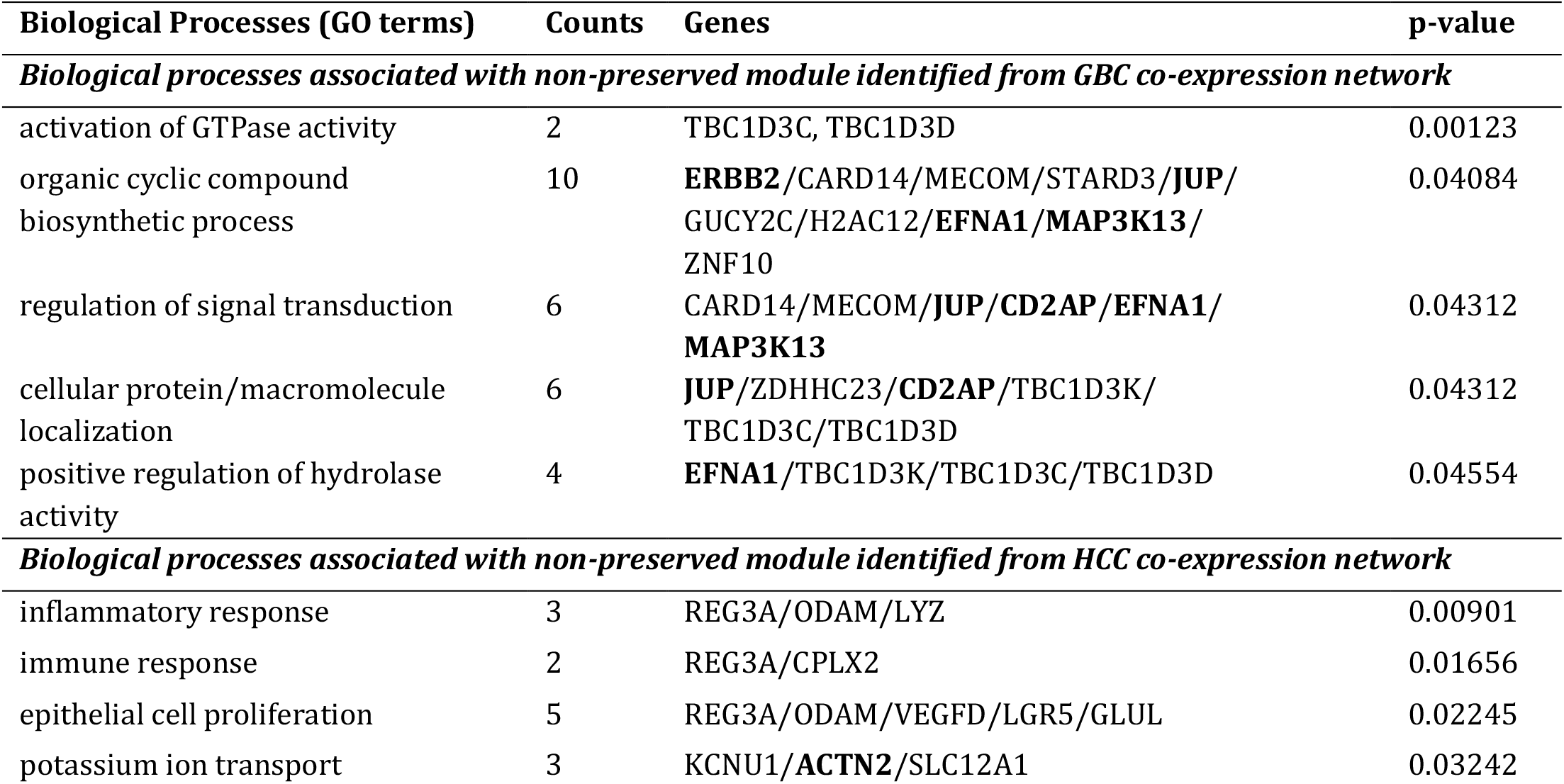
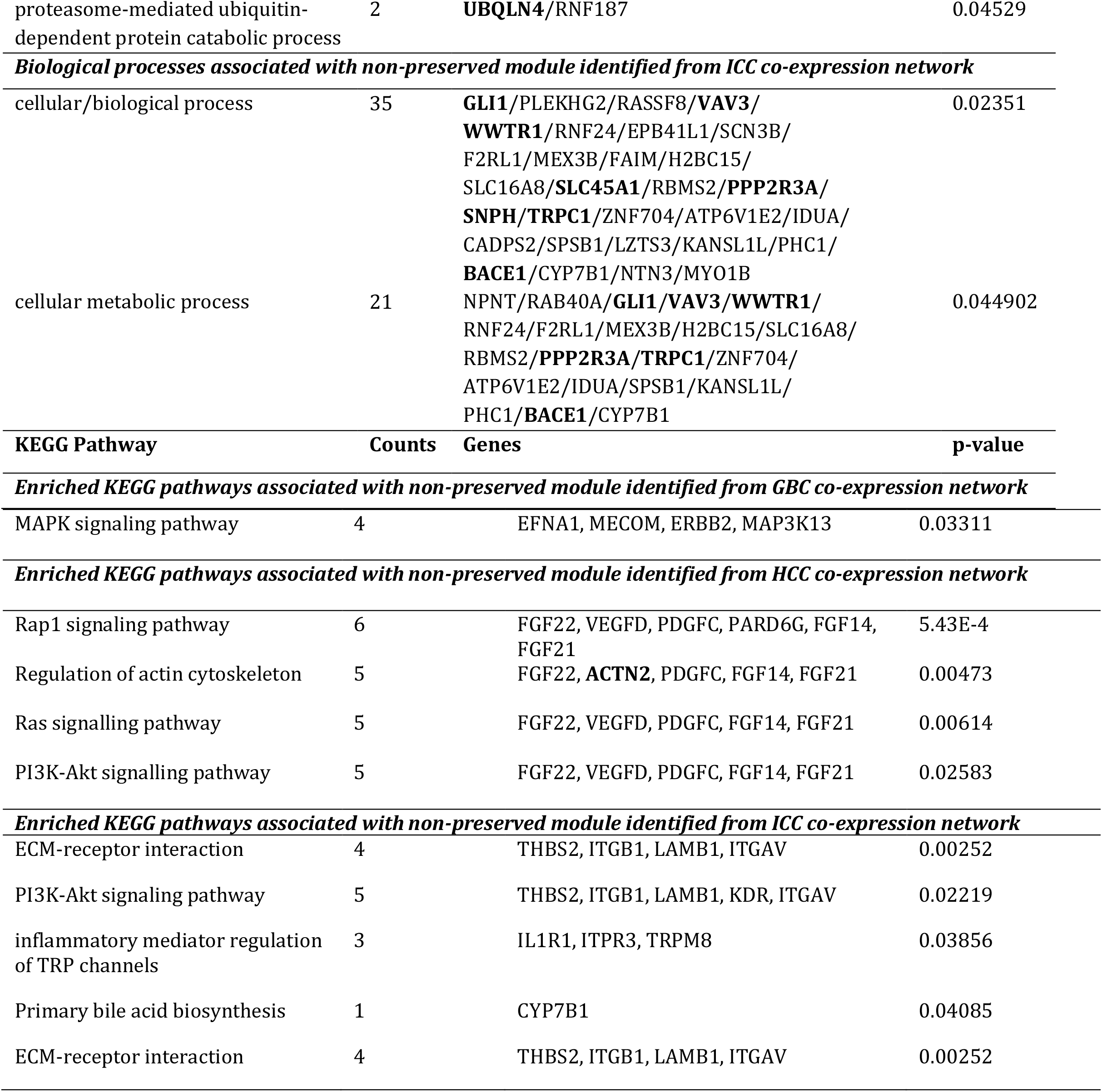
Significant biological processes and KEGG pathways associated with the non-preserved modules identified from GBC, HCC, and ICC network.

| Biological Processes (GO terms) | Counts | Genes | p-value |
| --- | --- | --- | --- |
| *Biological processes associated with non-preserved module identified from GBC co-expression network* |  |  |  |
| activation of GTPase activity | 2 | TBC1D3C, TBC1D3D | 0.00123 |
| organic cyclic compound biosynthetic process | 10 | **ERBB2**/CARD14/MECOM/STARD3/**JUP**/GUCY2C/H2AC12/**EFNA1**/**MAP3K13**/ZNF10 | 0.04084 |
| regulation of signal transduction | 6 | CARD14/MECOM/**JUP**/**CD2AP**/**EFNA1**/**MAP3K13** | 0.04312 |
| cellular protein/macromolecule localization | 6 | **JUP**/ZDHHC23/**CD2AP**/TBC1D3K/TBC1D3C/TBC1D3D | 0.04312 |
| positive regulation of hydrolase activity | 4 | **EFNA1**/TBC1D3K/TBC1D3C/TBC1D3D | 0.04554 |
| *Biological processes associated with non-preserved module identified from HCC co-expression network* |  |  |  |
| inflammatory response | 3 | REG3A/ODAM/LYZ | 0.00901 |
| immune response | 2 | REG3A/CPLX2 | 0.01656 |
| epithelial cell proliferation | 5 | REG3A/ODAM/VEGFD/LGR5/GLUL | 0.02245 |
| potassium ion transport | 3 | KCNU1/**ACTN2**/SLC12A1 | 0.03242 |
| proteasome-mediated ubiquitin-dependent protein catabolic process | 2 | UBQLN4/RNF187 | 0.04529 |
| **Biological processes associated with non-preserved module identified from ICC co-expression network** |  |  |  |
| cellular/biological process | 35 | GLI1/PLEKHG2/RASSF8/VAV3/<br>WWTR1/RNF24/EPB41L1/SCN3B/<br>F2RL1/MEX3B/FAIM/H2BC15/<br>SLC16A8/SLC45A1/RBMS2/PPP2R3A/<br>SNPH/TRPC1/ZNF704/ATP6V1E2/IDUA/<br>CADPS2/SPSB1/LZTS3/KANSL1L/PHC1/<br>BACE1/CYP7B1/NTN3/MYO1B | 0.02351 |
| cellular metabolic process | 21 | NPNT/RAB40A/GLI1/VAV3/WWTR1/<br>RNF24/F2RL1/MEX3B/H2BC15/SLC16A8/<br>RBMS2/PPP2R3A/TRPC1/ZNF704/<br>ATP6V1E2/IDUA/SPSB1/KANSL1L/<br>PHC1/BACE1/CYP7B1 | 0.044902 |
| **KEGG Pathway** | **Counts** | **Genes** | **p-value** |
| **Enriched KEGG pathways associated with non-preserved module identified from GBC co-expression network** |  |  |  |
| MAPK signaling pathway | 4 | EFNA1, MECOM, ERBB2, MAP3K13 | 0.03311 |
| **Enriched KEGG pathways associated with non-preserved module identified from HCC co-expression network** |  |  |  |
| Rap1 signaling pathway | 6 | FGF22, VEGFD, PDGFC, PARD6G, FGF14, FGF21 | 5.43E-4 |
| Regulation of actin cytoskeleton | 5 | FGF22, ACTN2, PDGFC, FGF14, FGF21 | 0.00473 |
| Ras signalling pathway | 5 | FGF22, VEGFD, PDGFC, FGF14, FGF21 | 0.00614 |
| PI3K-Akt signalling pathway | 5 | FGF22, VEGFD, PDGFC, FGF14, FGF21 | 0.02583 |
| **Enriched KEGG pathways associated with non-preserved module identified from ICC co-expression network** |  |  |  |
| ECM-receptor interaction | 4 | THBS2, ITGB1, LAMB1, ITGAV | 0.00252 |
| PI3K-Akt signaling pathway | 5 | THBS2, ITGB1, LAMB1, KDR, ITGAV | 0.02219 |
| inflammatory mediator regulation of TRP channels | 3 | IL1R1, ITPR3, TRPM8 | 0.03856 |
| Primary bile acid biosynthesis | 1 | CYP7B1 | 0.04085 |
| ECM-receptor interaction | 4 | THBS2, ITGB1, LAMB1, ITGAV | 0.00252 |

### 2.6 Analysis of transcription factors (TFs) regulatory network and identification of hub TFs

From the unique sDEGs identified for each cancer type, 21, 62, and 96 code for differentially expressed transcription factors in GBC, HCC, and ICC, respectively. TF-TG regulatory networks were constructed using the TFs as source nodes and their target genes as target nodes for each cancer as indicated in **Figure 5**. The topological analysis of the regulatory networks such as assortativity, path length is computed using igraph, an R package. The assortativity of GBC, HCC, and ICC network is negative i.e., −0.055, −0.059 and −0.079, respectively, which implies that nodes with higher degree interact with nodes having low degree centrality. The top 5 hub TFs identified for all the 3 cancers based on degree centrality have been tabulated in **supplementary Table 2**. From the TRNs of GBC, HCC, and ICC, it can be observed that the FOX families of TFs are conserved as hubs in all the cancer types, with ZNF family of proteins overlapping among GBC and ICC and KLF TFs common between ICC and HCC indicating the potential role of these TF families in development of hepatobiliary cancers.

**Figure 5:**
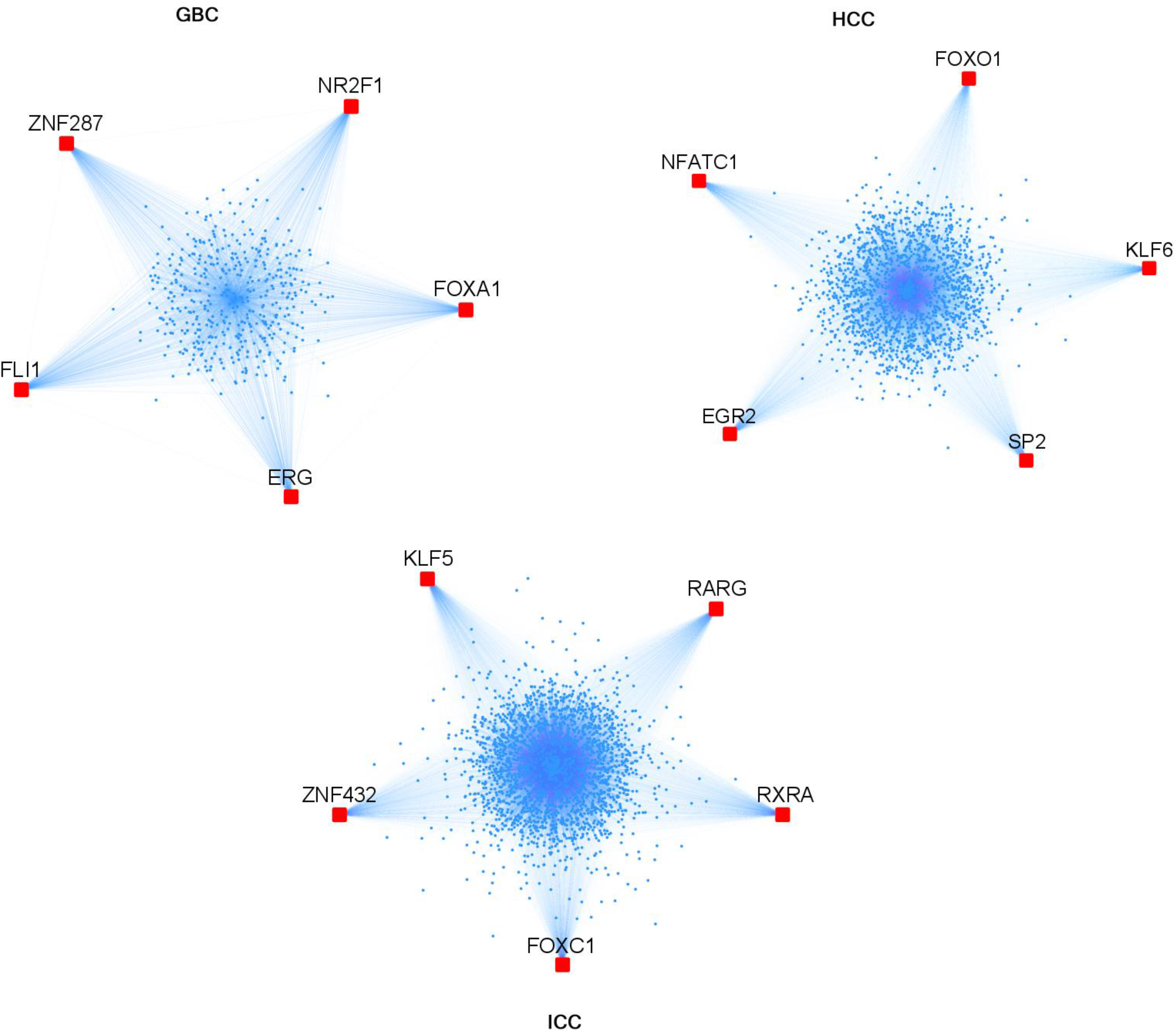
TF-TG network of the unique DEGs in GBC, HCC, and ICC. The red nodes represent the top hub TFs having the highest degree, and the small blue nodes represent target genes. The TF-TG interactions are identified through PWM scanning of 1kbp upstream sequences of sDEGs that assign a probability score of the transcription factor binding to the target gene.

### 2.7. Independent validation of the identified hub genes

The gene expression level of the potential hub genes and hub transcription factors are validated using the GEPIA database. The expression level of each hub gene has been represented as box plots (**Figure 6a**). From the boxplots, we observe that all the hub genes and TFs are overexpressed in HCC and CCA patients’ samples from TCGA-LIHC and TCGA-CCA dataset as compared to normal tissue samples (*p-*value < 0.01 and log fold change of 1). Since, there is no TCGA-GBC dataset in the GEPIA database; we validated the hub genes of GBC by taking TCGA-CCA dataset. It is observed that hub genes from GBC are significantly overexpressed in CCA samples from TCGA, which indicates that these hub genes might also be significantly upregulated in GBC as GBC and CCA are part of the same organ system. Furthermore, the cBioPortal has been used to evaluate the genomic aberrations of the hub genes by taking studies from GBC (MSK, Cancer 2018), HCC (TCGA, Firehouse Legacy), and CCA (TCGA, Firehouse Legacy). The OncoPrint tool of cBioPortal shows that 20% (101/502) of patients’ cases undergo genetic alterations such as amplification, deletion and several mutations (**Figure 6b**).

**Figure 6:**
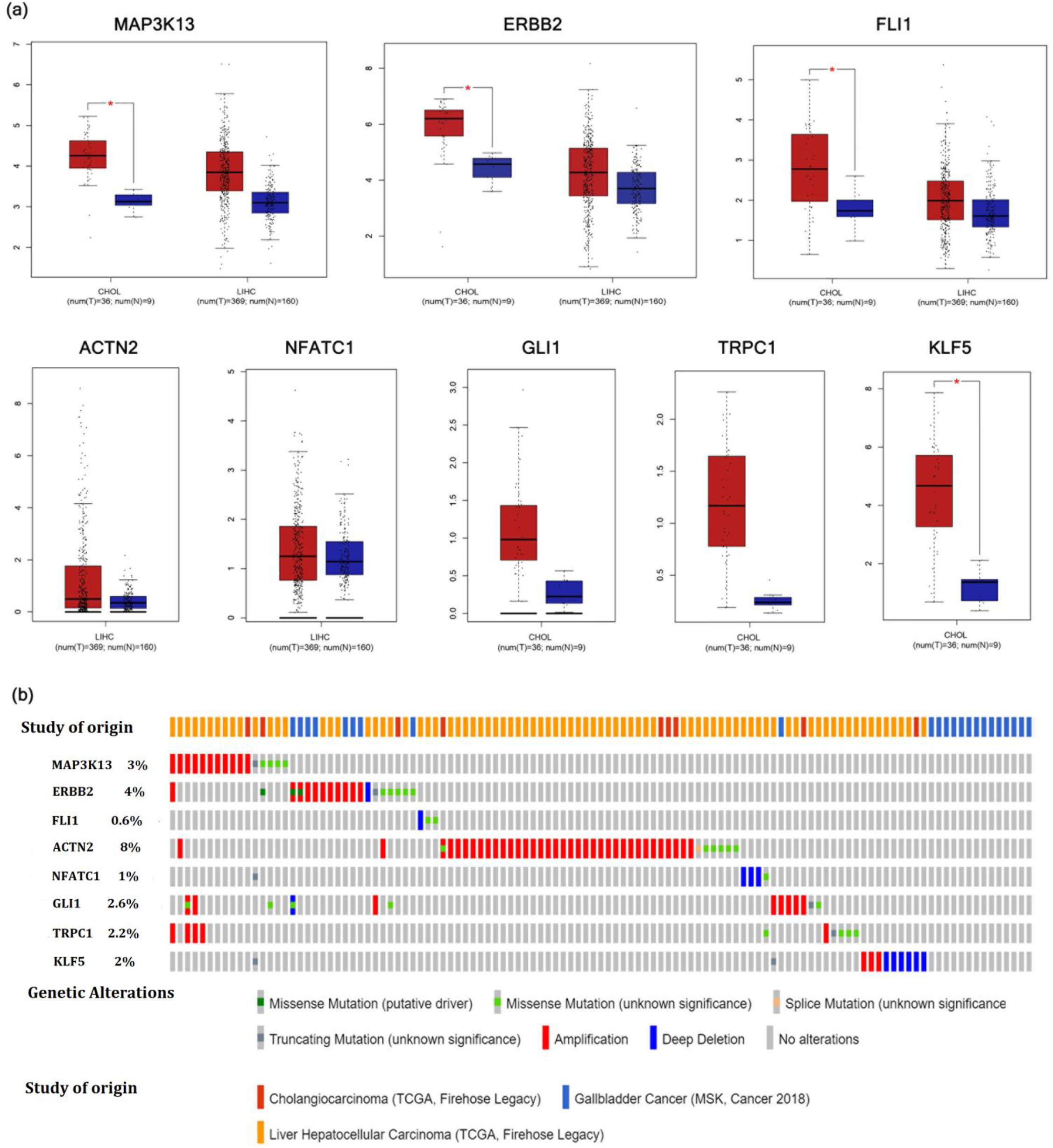
Validation of the expression of hub genes and hub TFs and detection of genetic alteration. The gene expression validation of the hub genes identified from GBC, HCC and ICC from respective TCGA datasets. The red and blue box represents tumor and normal samples respectively. (b) The genetic alterations associated with hub genes and hub TFs

## 3. Discussions

An effective systems biology approach called CUMSI has been developed and benchmarked, that enables the identification of unique and overlapping DEGs using publicly available transcriptomic datasets for GBC, HCC, and ICC. The 256 overlapping DEGs among the 3 cancers follow a differential pattern of gene expression and are mostly associated with cell cycle processes. In total 561, 2005, and 2580 unique DEGs have been identified for GBC, HCC, and ICC respectively. Using GSEA, the unique DEGs in GBC, HCC, and ICC were found to be associated with cellular processes, immune system regulation, and lipid metabolic processes, respectively. Using the unique DEGs, gene co-expression networks, transcription factor-target gene (TF-TG) networks, and protein-protein interaction (PPI) networks are constructed for each cancer type to identify the key players (hub genes) involved in carcinogenesis.

The hub genes identified for GBC through gene co-expression network analysis and PPI network analysis are found to be largely associated with MAPK signaling, G-protein coupled receptors, RET signaling, zinc dependent signaling, cytoskeleton proteins, stem cell differentiation, angiogenesis, and immune modulation. Mitogen activated signaling pathway has been implicated in many cancers, including GBC and mediates multiple cellular processes like cell differentiation, proliferation, apoptosis, and migration by phosphorylation of many downstream targets (Buchegger et al., 2017). MAP3K13, identified as a hub gene in GBC, has recently been reported to stabilize the proto-oncogene c-Myc. Direct inhibition of c-Myc has been unachievable till date and therefore, MAP3K13 presents a prospective therapeutic target for curbing oncogenesis (Q. Zhang et al., 2020). The MAPK signaling cascade can be dysregulated through aberrations in epidermal growth factor (EGFR) family of proteins known to upregulate the activation of pro-oncogenic signaling pathways like AKT-PI3K-mTOR, Ras-Raf-MEK-ERK-MEK MAPK pathway (Wee & Wang, 2017). ERBB2, a member of the EGFR family, has been identified as a hub protein in GBC. Recent study suggests that aberrations in ERBB2are associated with early-stage gallbladder tumors and therefore, can serve as a marker for early diagnosis and a therapeutic target for gallbladder tumors (Iyer et al., 2019). ERBB2 is also involved in RET signaling pathway that has been implicated in the development of the mammalian enteric nervous system (ENS) (Pachnis et al., 1998). The nervous system plays an integral role in the formation of the tumor microenvironment (TME) and has been recognized in the progression of gastrointestinal cancers. There is ongoing research on the use of denervation to inhibit tumorigenesis and can further be explored for its use in therapy (Schonkeren et al., 2021).

The hub genes identified for HCC largely comprise of pseudogenes, cytoskeletal proteins, DNA repair molecules, zinc finger proteins, genes for immune system regulation and ubiquitin proteins. Pseudogenes are largely regarded as non-functional or silent genes but, in recent years, they have emerged as highly specific biomarkers with potential use for diagnosis and prognosis of diseases. Most of the pseudogenes discovered till date are associated with human cancer, exerting parental gene related as well as unrelated functions and thus, evolving as key players in complex regulatory networks (Poliseno et al., 2015). In HCC, 3 pseudogenes are identified as hubs with AC069287.1 having the highest degree. The Ensembl database for *Homo sapiens* reports AC069287.1 as a long noncoding RNA gene. In addition to pseudogenes, HCC has been largely associated with immune signaling pathways with NFATC1, a key regulator of T-cell differentiation, identified as a hub gene. NFATC1 plays an important role in inducing the expression of cytokine genes IL-2 and IL-4 in T-cells (Zhao et al., 2010). A recent study has highlighted the role of NFATC1 in the activation of a tumor necrosis factor (TNF) receptor protein promoting tumor metastasis. This cascade can therefore, be targeted for developing anti-metastatic drugs for cancer patients (Liang et al., 2021).

The hub genes in ICC are associated with ion channel proteins (calcium), PI3-Akt signaling, zinc finger proteins, GTPases, stem cell differentiation pathways, aspartic proteases, bile acid biosynthesis, and retinoic acid signaling. TRPC1 and BACE1 are identified has hub genes through both gene co-expression network analysis and PPI network analysis. TRPC1 is a cation channel protein that plays a key role in maintaining Ca^2+^ balance in the cell. Ca^2+^serves as a secondary messenger for a myriad of physiological cellular processes such as cell proliferation, differentiation, apoptosis, migration, and metastasis. In some cancer types, TRPC1 has been associated with a poor prognosis whereas in others, it has been associated with a good clinical outcomes (Elzamzamy et al., 2020; Zeng et al., 2021). An in-depth study of TRPC1 with respect to clinical outcomes can provide further evidence for its use as a prognostic marker for ICC patients.BACE1 is an aspartic protease responsible for cleaving amyloid precursor protein (APP) to neurotoxic peptides implicated in Alzheimer’s disease. It has been suggested that BACE1 mediated APP maturation promotes the formation of neutrophil extracellular traps (NETs), a defense mechanism wherein neutrophils exude decondensed chromatin into the extracellular space, trapping tumor cells and promoting carcinogenesis. Further studies can help elucidate the molecular mechanisms through which BACE1 drives cancer progression and help explore its use as a potential therapeutic target (Farris et al., 2021).

Through TF-TG network analysis, we identify 21, 62, and 96 transcription factors to be differentially expressed in GBC, HCC, and ICC respectively. Zinc finger proteins (ZNF, KLF family) and forkhead box (FOX) transcription factors were found to be overlapping between the 3 cancers. FOX proteins are characterized by 80-100 amino acid DNA-binding motifs on the basis of which, the FOX family is classified into various subclasses. These proteins are largely involved in cell differentiation, proliferation, cell cycle control, and development. FOXA, FOXC, FOXM, FOXO, and FOXP subclasses have largely been implicated in tumorigenesis with FOXA1, FOXO1, and FOXC1 identified as hub TFs through our study in GBC, HCC, and ICC respectively (W. Zhang et al., 2017). Zinc finger proteins are characterized by a zinc finger structure maintained by zinc ions that coordinate cysteine and histidine residues. ZNFs have been associated with transcriptional regulation, ubiquitin-mediated protein degradation, signal transduction, actin targeting, DNA repair, and cell migration, thereby leading to tumorigenesis, cancer progression and metastasis formation. ZNFs may be acting as both oncogenes and/or tumor suppressor genes. Further studies into the molecular mechanisms of these proteins can provide substantial leads for diagnostic or therapeutic biomarkers (Cassandri et al., 2017).

The identified hub genes for each cancer are unique, and yet associated with overlapping as well as unique molecular signatures, which highlights the intricate association of the organs of the hepatobiliary system. Further in-depth study on these genes can provide leads to developing potential diagnostic, prognostic biomarkers and therapeutic targets.

In the present study, our approach of CUMSI enables us to identify common and unique molecular signatures, pathways, and processes associated with three aggressive cancers of the hepatobiliary system-GBC, HCC and ICC in a systems level perspective. The key processes and pathways identified with liver cancer progression and intrahepatic cholangiocarcinoma are associated with immune system regulation and lipid metabolic pathways respectively. Under normal physiological conditions, the liver is responsible for mediating immune tolerance, and the bile acids secreted into the bile ducts regulate lipid metabolism. This suggests that carcinogenesis results from the dysregulation of pathways involved in carrying out the primary function of that organ. Our study showed that the key genes and pathways identified for each individual cancer of the HBS are linked with the primary function of each organ and each cancer exhibits a unique expression pattern despite being part of the same system. This indicates that cancer pathogenesis and progression have unique molecular signatures. Therefore each cancer type requires strategies for targeted therapy. The potential hub genes, hub TFs and significant pathways identified during this analysis from individual cancers of the HBS may be potent diagnostic and prognostic biomarkers in patients with HBCs.

## 4. Materials and method

### 4.1 RNAseq dataset retrieval and analysis

To obtain the relevant benchmarked datasets on GBC, HCC, ICC and ECC for this study, a comprehensive search was conducted on the NCBI Gene Expression Omnibus database using the following criteria – (i) study type: expression profiling by high throughput sequencing, (ii) attribute name: tissue and (iii) organism: Homo sapiens. There were 5 GBC, 117 HCC, 9 ICC and 0 ECC datasets available on NCBI conforming to the above filters. For this study, datasets containing (i) paired end data, (ii) both case-control samples, (iii) sample size≥20, and (iv) information about the sequencing platform used as well as the experimental protocol selected for analysis. We considered 3 datasets – **GSE139682** for GBC, **GSE105130** for HCC and **GSE119336** for ICC (**Supplementary Table 1**). The datasets were downloaded from the ENA database in Fastq format, and the raw reads were filtered based on their quality score using FastQC. The processed reads were aligned with the reference human genome *Homo sapiens* (GRCh38) using Hisat2(Kim et al., 2015) and quantified at the gene level using featureCount to obtain the count matrix for each gene (Liao et al., 2014).

### 4.2 Differential gene expression analysis

After the generation of count data from transcriptomic data analysis, it is essential to perform pre-processing steps comprising of low read count removal, normalization, and transformation on the raw count data before downstream analysis. Otherwise, technical variability such as differences in library size (the total number of reads per sample) and library composition (the set of genes to which all reads of a sample are mapped) can falsely reflect differential gene expression (Chowdhury et al., 2020; Evans et al., 2018; Sahu et al., 2020). Widely used differential gene expression analysis (DGEA) tools-DESeq2 and EdgeR follow DESeq2 and TMM normalization methods respectively to account for these biases. The DESeq2 method calculates the ratio of each read count to the geometric (logarithmic) mean of all read counts for that gene across all samples. The median of these ratios is subsequently used for scaling each sample. The Trimmed Mean of M-values chooses a reference sample and calculates fold changes and absolute expression levels relative to that sample (Evans et al., 2018). These tools were used for the normalization and transformation of our datasets. Applying *padj≤0*.*05* and log2FC ±1, the final list of significant DEGs from DESeq2 and EdgeR were obtained. Consensus of both the DGEA tools was taken for obtaining common as well as unique DEGs for GBC, HCC, and ICC (Sahu et al., 2020).

### 4.3 Construction of gene co-expression network and screening of non-preserved modules

Gene co-expression networks are useful for finding correlations in gene expression patterns across samples. The WGCNA package available in R has been used for constructing co-expression networks for each cancer and their respective controls. Outlier samples were first removed by hierarchical clustering for computing the appropriate β value. The β value reflects the lowest integer to which the equation for estimating connection strengths between genes should be raised [Adj_ij_=(abs(cor(g_i_, g_j_)))^β^] for obtaining a scale free network. Such networks are desirable and consist of a small number of highly connected nodes called hubs. Hub genes play an important role in the molecular mechanisms determining the disease progression.

The main steps of WGCNA are – (i) computation of pairwise gene similarity using Pearson’s correlations to construct an adjacency matrix (Adj_ij_), (ii) formation of a scale free co-expression network from the adjacency matrix taking into account β value (soft thresholding parameter), (iii) identification of modules by deriving Topological Overlap Matrix (TOM) from adjacency matrix and perform hierarchical clustering and dynamic tree cut, (iv) distinguishing non-preserved modules using *Z-summary* and *medianRank* statistics and (v) performing gene set enrichment and pathway analysis for the non-preserved modules to identify hub genes. Preservation analysis is important to identify modules that are subjected to change across conditions, indicating that they play a role in pathogenesis. A well-preserved module (*Z-summary>10*), can be found in both the reference network and the test network, but a non-preserved module (*Z-summary<2*) or a weakly-preserved module (*2<Z-summary<10*) can only be found in either of the networks. A non-preserved module is distinguished by a low *Z-summary* score and a high *medianRank* that is subsequently used for identification of potential candidate genes (Langfelder & Horvath, 2008; Sahu et al., 2020).

### 4.4. Hub gene identification through intra-modular connectivity

In network biology, intra-modular connectivity describes how connected a gene is to all the other genes within a module. This is measured from the Topological Overlap Matrix (TOM) that takes into account, not only the co-expression information but also, the relative inter-connectedness or the shared neighbours between 2 nodes (Langfelder, 2013; B. Zhang & Horvath, 2005). Here, we analyzed the intramodular connectivity of the non-preserved modules for identifying hub genes for each cancer type of the HBS. PPI networks are constructed using the STRING database that predicts interaction of the genes at the protein level (Szklarczyk et al., 2019). Significant interactions are filtered using a combined score> 0.7 and PPI networks of the genes comprising the non-preserved modules are constructed using the NetworkAnalyst tool (Zhou et al., 2019). The genes having a high degree, i.e. a large number of neighbors/interacting proteins have been identified as hub proteins. These hubs may play crucial roles in GBC, HCC, and ICC pathogenesis (**Figure 2**).

### 4.5. Gene Regulatory Network (GRN) analysis

For the construction of the Gene Regulatory Network, 1KB upstream sequences of the unique DEGs (561, 2005, 2580) for GBC, HCC and ICC have been extracted using the RSAT (Regulatory Sequence Analysis Tools) database (Thomas-Chollier et al., 2008). PWMs (Position Weight Matrices) for all human TFs are obtained using the cis-BP database (Dimitrova et al., 2014). A PWM is a model reflecting the binding specificity of a TF and is used to scan the upstream sequences of the DEGs for establishing transcription factor-target gene interactions (Aerts, 2012). PWM scanning has been carried out using MEME suite (T. L. Bailey et al., 2009). The obtained TF-TG interactions are visualized using Cytoscapev3.8.2(Cline et al., 2007). The top 5 TFs are obtained for each cancer type (GBC, HCC, ICC) by sorting them based on degree and considered as hubs for subsequent analysis.

### 4.6. Gene expression validation of the hub genes and TFs

The Gene Expression Profiling Interactive Analysis (GEPIA) database has been used for validation of the expression of hub genes identified from WGCNA analysis and TRN analysis. GEPIA is a web server that gives customizable interactive analysis of the gene expression based on *The Cancer Genome Atlas* (TCGA) and GTEx data. For the validation of hub genes, we considered fold change value > 2 and *p-*value < 0.01 as statistically significant. We evaluated genetic alterations such as mutations and copy number alterations linked with the potential hub genes and TFs using cBioPortal database (https://www.cbioportal.org/). The results generated from cBioPortal were visualized as OncoPrint.

## Acknowledgement

Dr. Pankaj Barah would like to acknowledge Department of Biotechnology, Government of India for providing the Ramalingaswami Re-entry Fellowship grant.

## Author contribution

P.B and D.K.B contributed to the conceptualization and study design; P.B and D.K.B participated in the methodology; N.R and R.L analyzed the data; N.R and R.L contributed to the draft preparation. All the authors edited and approved the final version of the manuscript. All authors have read and agreed to the published version of the manuscript.

## Dataset availability

The datasets used for this study are available at NCBI-GEO database (GEO Accession No. GSE139682, GSE105130 and GSE119336).

## Supplementary Data

**Supplementary Figure S1:** sDEGs obtained through DESeq2+EdgeR consensus for GBC, HCC, and ICC.

**Supplementary Figure S2: Functional enrichment analysis**. Biological processes (left) & KEGG pathways (right) associated with unique DEGs of **(a)** GBC, **(b)** HCC,&**(c)** ICC.

**Supplementary Figure S3: Overlapping sDEG analysis. (a)** Heatmap constructed using log2FC values of 256 overlapping sDEGs between GBC, HCC, and ICC. **(b)** Significant biological processes associated with overlapping sDEGs.

**Supplementary Figure S4:** β power value for cancer (left) and control (right) gene co-expression network construction for GBC, HCC, and ICC.

**Supplementary Tables**

**Table S1:** Details of 3 datasets – GSE139682 for GBC, GSE105130 for HCC and GSE119336 for ICC downloaded from the ENA database in Fastq format,

**Table S2:** *Z-summary* preservation of modules in GBC, HCC, and ICC network.

**Table S3:** Identification of hub TFs through TF-TG network analysis

## References

Aerts, S. (2012). Computational Strategies for the Genome-Wide Identification of cis-Regulatory Elements and Transcriptional Targets. In Current Topics in Developmental Biology (1st ed., Vol. 98). Elsevier Inc. https://doi.org/10.1016/B978-0-12-386499-4.00005-7

Bailey, A., & Shah, S. A. (2019). Screening high risk populations for cancer: Hepatobiliary. Journal of Surgical Oncology, 120(5), 847–850. https://doi.org/10.1002/jso.25633

Bailey, T. L., Boden, M., Buske, F. A., Frith, M., Grant, C. E., Clementi, L., Ren, J., Li, W. W., & Noble, W. S. (2009). MEME Suite: Tools for motif discovery and searching. Nucleic Acids Research, 37(SUPPL. 2), 202–208. https://doi.org/10.1093/nar/gkp335

Brägelmann, J., Barahona Ponce, C., Marcelain, K., Roessler, S., Goeppert, B., Gallegos, I., Colombo, A., Sanhueza, V., Morales, E., Rivera, M. T., de Toro, G., Ortega, A., Müller, B., Gabler, F., Scherer, D., Waldenberger, M., Reischl, E., Boekstegers, F., Garate-Calderon, V., … Lorenzo Bermejo, J. (2021). Epigenome-Wide Analysis of Methylation Changes in the Sequence of Gallstone Disease, Dysplasia, and Gallbladder Cancer. In Hepatology (Vol. 73, Issue 6, pp. 2293–2310). https://doi.org/10.1002/hep.31585

Buchegger, K., Silva, R., López, J., Ili, C., Araya, J. C., Leal, P., Brebi, P., Riquelme, I., & Roa, J. C. (2017). The ERK/MAPK pathway is overexpressed and activated in gallbladder cancer. Pathology Research and Practice, 213(5), 476–482. https://doi.org/10.1016/j.prp.2017.01.025

Cassandri, M., Smirnov, A., Novelli, F., Pitolli, C., Agostini, M., Malewicz, M., Melino, G., & Raschellà, G. (2017). Zinc-finger proteins in health and disease. Cell Death Discovery, 3(1). https://doi.org/10.1038/cddiscovery.2017.71

Chowdhury, H. A., Bhattacharyya, D. K., & Kalita, J. K. (2020). (Differential) Co-Expression Analysis of Gene Expression: A Survey of Best Practices. IEEE/ACM Transactions on Computational Biology and Bioinformatics, 17(4), 1154–1173. https://doi.org/10.1109/TCBB.2019.2893170

Cline, M. S., Smoot, M., Cerami, E., Kuchinsky, A., Landys, N., Workman, C., Christmas, R., Avila-Campilo, I., Creech, M., Gross, B., Hanspers, K., Isserlin, R., Kelley, R., Killcoyne, S., Lotia, S., Maere, S., Morris, J., Ono, K., Pavlovic, V., … Bader, G. D. (2007). Integration of biological networks and gene expression data using cytoscape. Nature Protocols, 2(10), 2366–2382. https://doi.org/10.1038/nprot.2007.324

Craig, A. J., von Felden, J., Garcia-Lezana, T., Sarcognato, S., & Villanueva, A. (2020). Tumour evolution in hepatocellular carcinoma. Nature Reviews Gastroenterology and Hepatology, 17(3), 139–152. https://doi.org/10.1038/s41575-019-0229-4

Dimitrova, N., Zamudio, J. R., Jong, R. M., Soukup, D., Resnick, R., Sarma, K., Ward, A. J., Raj, A., Lee, J., Sharp, P. A., & Jacks, T. (2014). Determination and Inference of Eukaryotic Transcription Factor Sequence Specificity. Cell, 158(6), 1431–1443. https://doi.org/10.1016/j.cell.2014.08.009.Determination

Elzamzamy, O. M., Penner, R., & Hazlehurst, L. A. (2020). The Role of TRPC1 in Modulating Cancer Progression. Cells, 9(2), 1–13. https://doi.org/10.3390/cells9020388

Evans, C., Hardin, J., & Stoebel, D. M. (2018). Selecting between-sample RNA-Seq normalization methods from the perspective of their assumptions. Briefings in Bioinformatics, 19(5), 776–792. https://doi.org/10.1093/bib/bbx008

Farris, F., Matafora, V., & Bachi, A. (2021). The emerging role of β-secretases in cancer. Journal of Experimental and Clinical Cancer Research, 40(1), 1–10. https://doi.org/10.1186/s13046-021-01953-3

Giudicessi, J. R., & Ackerman, M. J. (2013). Determinants of incomplete penetrance and variable expressivity in heritable cardiac arrhythmia syndromes. Translational Research, 161(1), 1–14. https://doi.org/10.1016/j.trsl.2012.08.005

Iyer, P., Shrikhande, S. V., Ranjan, M., Joshi, A., Gardi, N., Prasad, R., Dharavath, B., Thorat, R., Salunkhe, S., Sahoo, B., Chandrani, P., Kore, H., Mohanty, B., Chaudhari, V., Choughule, A., Kawle, D., Chaudhari, P., Ingle, A., Banavali, S., … Dutt, A. (2019). ERBB2 and KRAS alterations mediate response to EGFR inhibitors in early stage gallbladder cancer. International Journal of Cancer, 144(8), 2008–2019. https://doi.org/10.1002/ijc.31916

Jusakul, A., Cutcutache, I., Yong, C. H., Lim, J. Q., Huang, M. N., Padmanabhan, N., Nellore, V., Kongpetch, S., Ng, A. W. T., Ng, L. M., Choo, S. P., Myint, S. S., Thanan, R., Nagarajan, S., Lim, W. K., Ng, C. C. Y., Boot, A., Liu, M., Ong, C. K., … Tan, P. (2017). Whole-genome and epigenomic landscapes of etiologically distinct subtypes of cholangiocarcinoma. Cancer Discovery, 7(10), 1116–1135. https://doi.org/10.1158/2159-8290.CD-17-0368

Kim, D., Langmead, B., & Salzberg, S. L. (2015). HISAT: A fast spliced aligner with low memory requirements. Nature Methods, 12(4), 357–360. https://doi.org/10.1038/nmeth.3317

Langfelder, P. (2013). Signed vs. Unsigned Topological Overlap Matrix. Technical Report, 3–4. https://labs.genetics.ucla.edu/horvath/CoexpressionNetwork/Rpackages/WGCNA/TechnicalReports/signedTOM.pdf

Langfelder, P., & Horvath, S. (2008). WGCNA: An R package for weighted correlation network analysis. BMC Bioinformatics, 9. https://doi.org/10.1186/1471-2105-9-559

Lazcano-Ponce, E. C., Miquel, J. F., Munoz, N., Herrero, R., Ferrecio, C., Wistuba, I. I., Alonso de Ruiz, P., Aristi Urista, G., & Nervi, F. (2001). Epidemiology and Molecular Pathology of Gallbladder Cancer. CA: A Cancer Journal for Clinicians, 51(6), 349–364. https://doi.org/10.3322/canjclin.51.6.349

Liang, Q., Wang, Y., Lu, Y., Zhu, Q., Xie, W., Tang, N., Huang, L., An, T., Zhang, D., Yan, A., Liu, S., Ye, L., & Zhu, C. (2021). RANK promotes colorectal cancer migration and invasion by activating the Ca2+-calcineurin/NFATC1-ACP5 axis. Cell Death and Disease, 12(4). https://doi.org/10.1038/s41419-021-03642-7

Liao, Y., Smyth, G. K., & Shi, W. (2014). FeatureCounts: An efficient general purpose program for assigning sequence reads to genomic features. Bioinformatics, 30(7), 923–930. https://doi.org/10.1093/bioinformatics/btt656

Mardinoglu, A., Boren, J., Smith, U., Uhlen, M., & Nielsen, J. (2018). Systems biology in hepatology: Approaches and applications. Nature Reviews Gastroenterology and Hepatology, 15(6), 365–377. https://doi.org/10.1038/s41575-018-0007-8

Nakamura, H., Arai, Y., Totoki, Y., Shirota, T., Elzawahry, A., Kato, M., Hama, N., Hosoda, F., Urushidate, T., Ohashi, S., Hiraoka, N., Ojima, H., Shimada, K., Okusaka, T., Kosuge, T., Miyagawa, S., & Shibata, T. (2015). Genomic spectra of biliary tract cancer. Nature Genetics, 47(9), 1003–1010. https://doi.org/10.1038/ng.3375

Nault, J. C., Calderaro, J., Di Tommaso, L., Balabaud, C., Zafrani, E. S., Bioulac-Sage, P., Roncalli, M., & Zucman-Rossi, J. (2014). Telomerase reverse transcriptase promoter mutation is an early somatic genetic alteration in the transformation of premalignant nodules in hepatocellular carcinoma on cirrhosis. Hepatology, 60(6), 1983–1992. https://doi.org/10.1002/hep.27372

Nepal, C., Zhu, B., O’Rourke, C. J., Bhatt, D. K., Lee, D., Song, L., Wang, D., Van Dyke, A. L., Choo-Wosoba, H., Liu, Z., Hildesheim, A., Goldstein, A. M., Dean, M., LaFuente-Barquero, J., Lawrence, S., Mutreja, K., Olanich, M. E., Lorenzo Bermejo, J., Ferreccio, C., … Koshiol, J. (2021). Integrative molecular characterisation of gallbladder cancer reveals micro-environment-associated subtypes. Journal of Hepatology, 74(5), 1132–1144. https://doi.org/10.1016/j.jhep.2020.11.033

Pachnis, V., Durbec, P., Taraviras, S., Grigoriou, M., & Natarajan, D. (1998). Neural injury, repair, and adaptation in the GI tract: III. Role of the RET signal transduction pathway in development of the mammalian enteric nervous system. American Journal of Physiology - Gastrointestinal and Liver Physiology, 275(2 38-2), 193–196. https://doi.org/10.1152/ajpgi.1998.275.2.g183

Poliseno, L., Marranci, A., & Pandolfi, P. P. (2015). Pseudogenes in human cancer. Frontiers of Medicine, 2(September), 1–8. https://doi.org/10.3389/fmed.2015.00068

Sahu, A., Chowdhury, H. A., Gaikwad, M., Chongtham, C., Talukdar, U., Phukan, J. K., Bhattacharyya, D. K., & Barah, P. (2020). Integrative network analysis identifies differential regulation of neuroimmune system in Schizophrenia and Bipolar disorder. Brain, Behavior, & Immunity - Health, 2(December 2019), 100023. https://doi.org/10.1016/j.bbih.2019.100023

Schonkeren, S. L., Thijssen, M. S., Vaes, N., Boesmans, W., & Melotte, V. (2021). The emerging role of nerves and glia in colorectal cancer. Cancers, 13(1), 1–13. https://doi.org/10.3390/cancers13010152

Shibata, T., Arai, Y., & Totoki, Y. (2018). Molecular genomic landscapes of hepatobiliary cancer. Cancer Science, 109(5), 1282–1291. https://doi.org/10.1111/cas.13582

Szklarczyk, D., Gable, A. L., Lyon, D., Junge, A., Wyder, S., Huerta-Cepas, J., Simonovic, M., Doncheva, N. T., Morris, J. H., Bork, P., Jensen, L. J., & Von Mering, C. (2019). STRING v11: Protein-protein association networks with increased coverage, supporting functional discovery in genome-wide experimental datasets. Nucleic Acids Research, 47(D1), D607–D613. https://doi.org/10.1093/nar/gky1131

Teufel, A. (2015). Bioinformatics and database resources in hepatology. Journal of Hepatology, 62(3), 712–719. https://doi.org/10.1016/j.jhep.2014.10.036

Teufel, A., Marquardt, J. U., & Galle, P. R. (2012). Novel insights in the genetics of HCC recurrence and advances in transcriptomic data integration. Journal of Hepatology, 56(1), 279–281. https://doi.org/10.1016/j.jhep.2011.05.035

Thomas-Chollier, M., Sand, O., Turatsinze, J. V., Janky, R., Defrance, M., Vervisch, E., Brohée, S., & van Helden, J. (2008). RSAT: regulatory sequence analysis tools. Nucleic Acids Research, 36(Web Server issue). https://doi.org/10.1093/nar/gkn304

Totoki, Y., Tatsuno, K., Covington, K. R., Ueda, H., Creighton, C. J., Kato, M., Tsuji, S., Donehower, L. A., Slagle, B. L., Nakamura, H., Yamamoto, S., Shinbrot, E., Hama, N., Lehmkuhl, M., Hosoda, F., Arai, Y., Walker, K., Dahdouli, M., Gotoh, K., … Shibata, T. (2014). Trans-ancestry mutational landscape of hepatocellular carcinoma genomes. Nature Genetics, 46(12), 1267–1273. https://doi.org/10.1038/ng.3126

Wee, P., & Wang, Z. (2017). Epidermal growth factor receptor cell proliferation signaling pathways. Cancers, 9(5), 1–45. https://doi.org/10.3390/cancers9050052

Xue, F., Yang, L., Dai, B., Xue, H., Zhang, L., Ge, R., & Sun, Y. (2020). Bioinformatics profiling identifies seven immune-related risk signatures for hepatocellular carcinoma. PeerJ, 2020(5), 1–19. https://doi.org/10.7717/peerj.8301

Yang, J. D., Hainaut, P., Gores, G. J., Amadou, A., Plymoth, A., & Roberts, L. R. (2019). A global view of hepatocellular carcinoma: trends, risk, prevention and management. Nature Reviews Gastroenterology and Hepatology, 16(10), 589–604. https://doi.org/10.1038/s41575-019-0186-y

Zeng, Y. Z., Zhang, Y. Q., Chen, J. Y., Zhang, L. Y., Gao, W. L., Lin, X. Q., Huang, S. M., Zhang, F., & Wei, X. L. (2021). TRPC1 Inhibits Cell Proliferation/Invasion and Is Predictive of a Better Prognosis of Esophageal Squamous Cell Carcinoma. Frontiers in Oncology, 11(March), 1–13. https://doi.org/10.3389/fonc.2021.627713

Zhang, B., & Horvath, S. (2005). A general framework for weighted gene co-expression network analysis. Statistical Applications in Genetics and Molecular Biology, 4(1). https://doi.org/10.2202/1544-6115.1128

Zhang, Q., Li, X., Cui, K., Liu, C., Wu, M., Prochownik, E. V., & Li, Y. (2020). The MAP3K13-TRIM25-FBXW7α axis affects c-Myc protein stability and tumor development. Cell Death and Differentiation, 27(2), 420–433. https://doi.org/10.1038/s41418-019-0363-0

Zhang, W., Duan, N., Song, T., Li, Z., Zhang, C., & Chen, X. (2017). The emerging roles of forkhead box (FOX) proteins in osteosarcoma. Journal of Cancer, 8(9), 1619–1628. https://doi.org/10.7150/jca.18778

Zhao, Q., Wang, X., Liu, Y., He, A., & Jia, R. (2010). NFATc1: Functions in osteoclasts. International Journal of Biochemistry and Cell Biology, 42(5), 576–579. https://doi.org/10.1016/j.biocel.2009.12.018

Zhou, G., Soufan, O., Ewald, J., Hancock, R. E. W., Basu, N., & Xia, J. (2019). NetworkAnalyst 3.0: A visual analytics platform for comprehensive gene expression profiling and meta-analysis. Nucleic Acids Research, 47(W1), W234–W241. https://doi.org/10.1093/nar/gkz240

